# Olfactory transmucosal SARS-CoV-2 invasion as port of Central Nervous System entry in COVID-19 patients

**DOI:** 10.1101/2020.06.04.135012

**Authors:** Jenny Meinhardt, Josefine Radke, Carsten Dittmayer, Ronja Mothes, Jonas Franz, Michael Laue, Julia Schneider, Sebastian Brünink, Olga Hassan, Werner Stenzel, Marc Windgassen, Larissa Rößler, Hans-Hilmar Goebel, Hubert Martin, Andreas Nitsche, Walter J. Schulz-Schaeffer, Samy Hakroush, Martin S. Winkler, Björn Tampe, Sefer Elezkurtaj, David Horst, Lars Oesterhelweg, Michael Tsokos, Barbara Ingold Heppner, Christine Stadelmann, Christian Drosten, Victor Max Corman, Helena Radbruch, Frank L. Heppner

## Abstract

The newly identified severe acute respiratory syndrome coronavirus 2 (SARS-CoV-2) causes COVID-19, a pandemic respiratory disease presenting with fever, cough, and often pneumonia. Moreover, thromboembolic events throughout the body including the central nervous system (CNS) have been described. Given first indication for viral RNA presence in the brain and cerebrospinal fluid and in light of neurological symptoms in a large majority of COVID-19 patients, SARS-CoV-2-penetrance of the CNS is likely. By precisely investigating and anatomically mapping oro- and pharyngeal regions and brains of 32 patients dying from COVID-19, we not only describe CNS infarction due to cerebral thromboembolism, but also demonstrate SARS-CoV-2 neurotropism. SARS-CoV-2 enters the nervous system via trespassing the neuro-mucosal interface in the olfactory mucosa by exploiting the close vicinity of olfactory mucosal and nervous tissue including delicate olfactory and sensitive nerve endings. Subsequently, SARS-CoV-2 follows defined neuroanatomical structures, penetrating defined neuroanatomical areas, including the primary respiratory and cardiovascular control center in the medulla oblongata.

## Introduction

There is increasing evidence that the SARS-CoV-2 not only affects the respiratory tract but also impacts the CNS resulting in neurological symptoms such as loss of smell and taste, headache, fatigue, nausea and vomiting in more than one third of COVID-19 patients^1,2^. Moreover, acute cerebrovascular diseases and impaired consciousness have been described^3^. While recent studies describe the presence of viral RNA in the brain and cerebrospinal fluid (CSF)^4,5^ but lack a prove for genuine SARS-CoV-2 manifestation, a systematic analysis of COVID-19 autopsy brains aimed at understanding the port of SARS-CoV-2 entry and distribution within the CNS is lacking^6^.

The neuroinvasive potential of evolutionarily-related coronaviruses (CoVs) such as SARS-CoV and MERS-CoV has previously been described^7–9^. SARS-CoV including SARS-CoV-2 enter human host cells primarily by binding to the cellular receptor angiotensin-converting enzyme 2 (ACE2) and by the action of the serine protease TMPRSS2 for S protein priming^10^. Supporting evidence comes from animal studies demonstrating that SARS-CoV is capable of entering the brain upon intranasal infection of mice expressing human ACE2^8,11^. However, it is not known, which cells in the olfactory mucosa express these molecules in steady state or under inflammatory or septic conditions^11^, while there is first evidence for ACE2 expression in neuronal and glial cells in the CNS^13,14^. Along that line it is of note, that the immunoglobulin superfamily member CD147, which is expressed in neuronal and non-neuronal cells in the CNS^15,16^, has been shown to act as alternative cellular port for SARS-CoV-2 invasion^17^. To gain a better understanding of SARS-CoV-2 neurotropism and its mechanism of CNS entry and distribution, we analyzed the cellular mucosal-nervous micro-milieu as first site of viral infection and replication, followed by a thorough regional mapping of the consecutive olfactory nervous tracts and defined CNS regions in 32 COVID-19 autopsy cases.

## Results

Out of 32 COVID-19 autopsy cases, either proven to be RT-qPCR-positive for SARS-CoV-2 prior to death (N=29 of 32), or with clinical presentation highly suggestive of COVID-19 (N=3 of 32), four patients (corresponding to 13%) presented with acute infarction due to ischemia caused by (micro)thrombotic/thromboembolic events within the CNS (Supplementary Table 1). Similarly, microthrombotic events were also detectable in the olfactory mucosa (Supplementary Figure 1).

Assessment of viral load by means of RT-qPCR in regionally well-defined tissue samples including olfactory mucosa (region (R1), olfactory bulb (R2), oral mucosa (uvula; R3), trigeminal ganglion (R4) and medulla oblongata (R5) demonstrated highest levels of SARS-CoV-2 copies per cell within the olfactory mucosa sampled directly beneath the cribriform plate (N=13 of 22; Figure 1). Lower levels of viral RNA were found in the cornea, conjunctiva and oral mucosa, highlighting the oral and ophthalmic routes as additional potential sites of SARS-CoV-2 CNS entry (Figure 1). Only in few cases the cerebellum was also positive (N=2 of 21), while the wall of the internal carotid artery, which served as an internal negative control, was found to be negative in all investigated cases (N=10). The assessment of subgenomic (sg) RNA as surrogate for active virus replication yielded a positive result in 4 out of 13 SARS-CoV-2 RNA-positive olfactory mucosa samples and in 2 out of 6 SARS-CoV-2 RNA-positive uvulae, but in none of the other tissues analyzed in this study (Supplementary Table 1). Patients with shorter disease duration were more likely to be tested positive for viral RNA in the CNS tissue (Supplementary Table 1). The anatomical proximity between neurons, nerve fibers and mucosa within the oro-and nasopharynx (Figure 2) and the reported clinical-neurological signs related to alteration in smell and taste perception suggest that SARS-CoV-2 exploits this neuro-mucosal interface as port of CNS entry. On the apical side of the olfactory mucosa, dendrites of olfactory receptor neurons (ORNs) project into the nasal cavity, while on the basal side axons of olfactory receptor neurons merge into fila, which protrude through the cribriform plate directly into the olfactory bulb (Figure 2), thereby also having contact with the CSF^18^.

**Figure 1:**
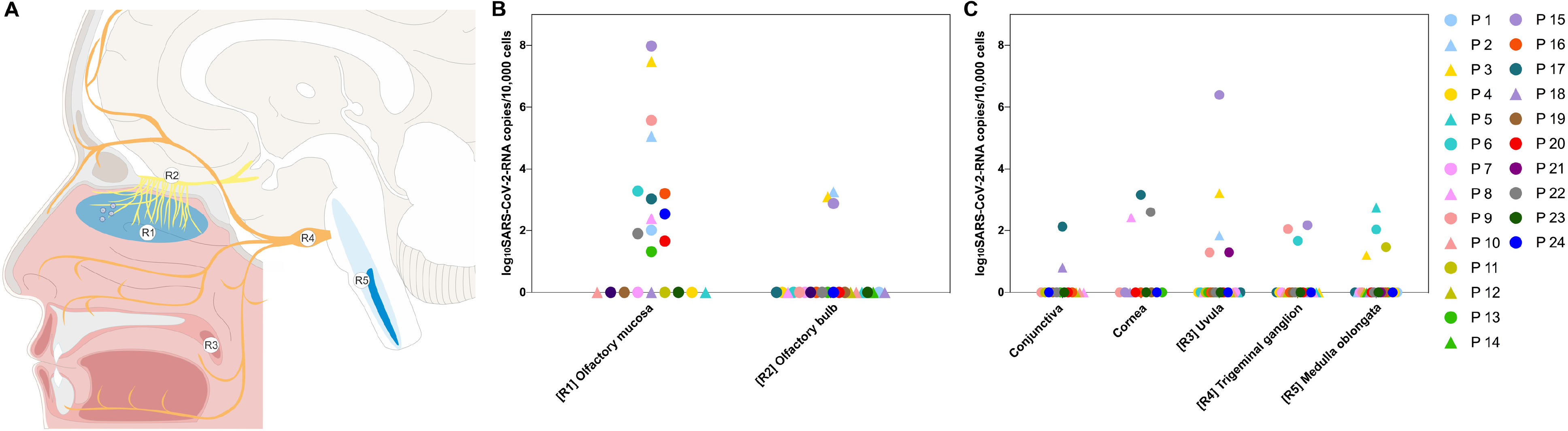
SARS-CoV-2 RNA levels of deceased COVID-19 patients in anatomically distinctly mapped oro- and nasopharyngeal as well as CNS regions. Cartoon depicting the anatomical structures sampled for histomorphological, ultrastructural and molecular analyses including SARS-CoV-2 RNA measurement from fresh (i.e. non-formalin-fixed) specimens of deceased COVID-19 patients (A). Specimens were taken from the olfactory mucosa underneath of the cribriform plate (Region (R)1), blue, N=22), the olfactory bulb (R2, yellow, N=23), from different branches of the trigeminal nerve (including conjunctiva (N=15), cornea (N=12), mucosa covering the uvula (R3, N=20)), the respective trigeminal ganglion in orange (R4, N=20), and the cranial nerve nuclei in the medulla oblongata (R5, dark blue, N=23). The quantitative results for each patient are shown in a logarithmic scale normalized on 10,000 cells (B, C). Female patients are displayed in triangular, male in circular symbols. P=patient.

**Figure 2:**
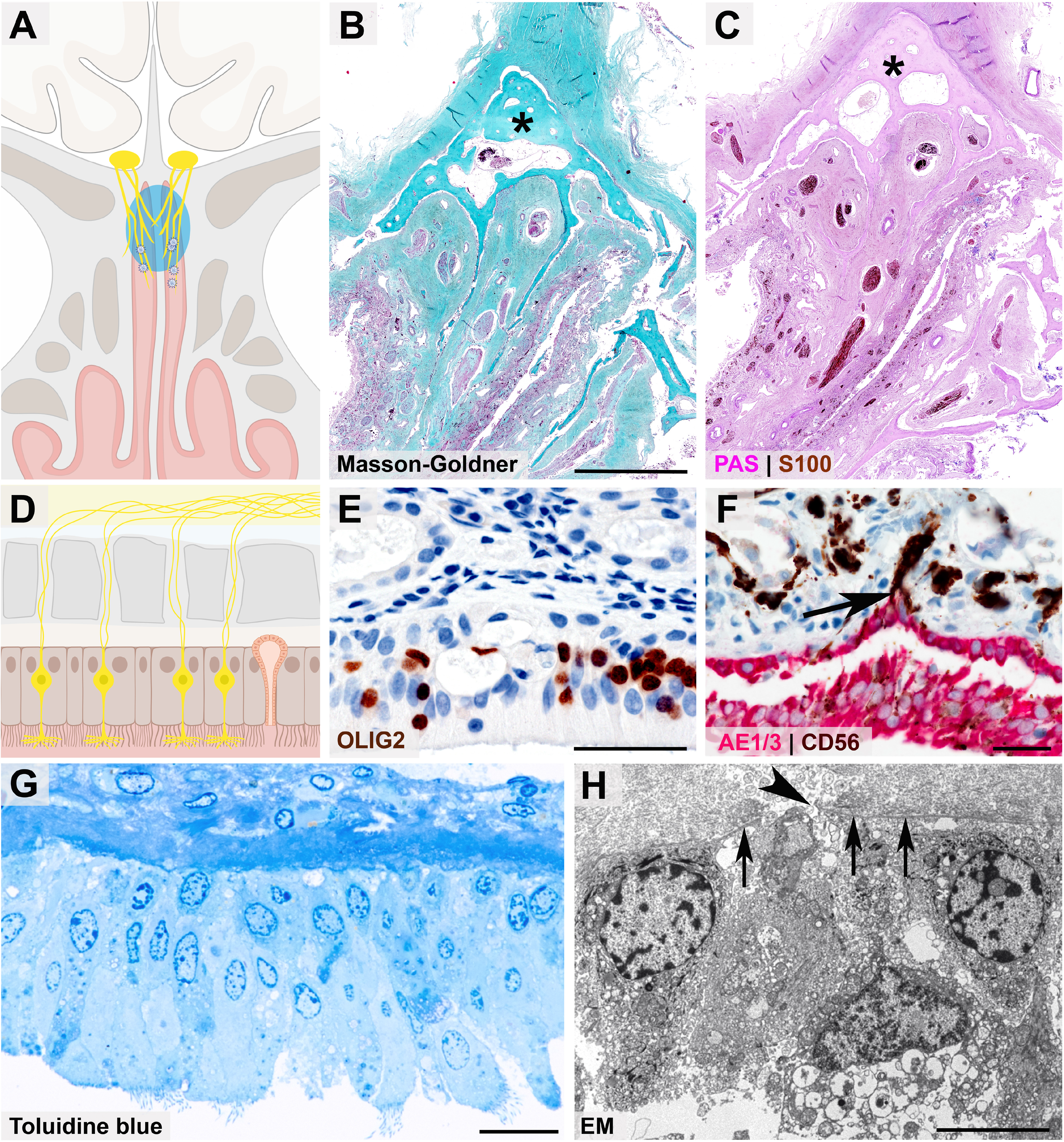
Close anatomical proximity of nervous and epithelial tissues in the olfactory mucosa. Cartoon (A) and histopathological coronal cross-sections (B - C) depicting the paranasal sinus region with the osseous cribriform plate (turquoise color and asterisk in B, pink color and asterisk in C) and the close anatomical proximity of the olfactory mucosa (green in B, purple in C) and nervous tissue characterized by nerve fibers immunoreactive for S100 protein (C, brown color). Cartoon (D) resembling the olfactory mucosa, which is composed of pseudostratified ciliated columnar epithelium, basement membrane, and lamina propria, also containing mucus-secreting Bowman glands and bipolar olfactory receptor neurons (ORNs), which coalesce the epithelial layer. Immunohistochemical staining of the olfactory mucosa (E, F) showing nuclear expression of OLIG2 specifying late neuronal progenitor and newly formed neurons (E, brown color)^31^, which are closely intermingled with epithelial cells (F, immunoreactivity for the pancytokeratin marker AE1/3, red color). The basement membrane underneath the columnar AE1/3-positive epithelium (F, red color) is discontinued due to CD56-positive (F, brown color) axonal projections of ORNs (F, arrow). The ORN dendrite carries multiple cilia and protrudes into the nasal cavity (G, semithin section, toluidine blue staining), while the axon (H, arrowhead) crosses the olfactory mucosa basement membrane (H, arrows) as a precondition for ORN projection into the glomeruli of the olfactory bulb, which is readily visible at the ultrastructural level). Scale bars: B: 3.5 cm; E, F: 50 μm; G: 20 μm; H: 5 μm.

Further evidence and support for a site-specific infection and inflammation by SARS-CoV-2 was provided by immunohistochemistry (Figure 3). Cells of the olfactory mucosa showed strong immunoreactivity in a characteristic perinuclear pattern when an antibody against the SARS-CoV spike protein was used. Furthermore, early activated macrophages formed small cell clusters in the epithelium expressing myeloid-related protein 14 (MRP14) (Supplementary Figure 2), initiating and regulating an immune cascade, which e.g. upon influenza virus infection, has been shown to orchestrate virus-associated inflammation by acting as endogenous damage-associated molecular pattern (DAMP), ultimately initiating TLR4-MyD88 signalling^19^.

**Figure 3:**
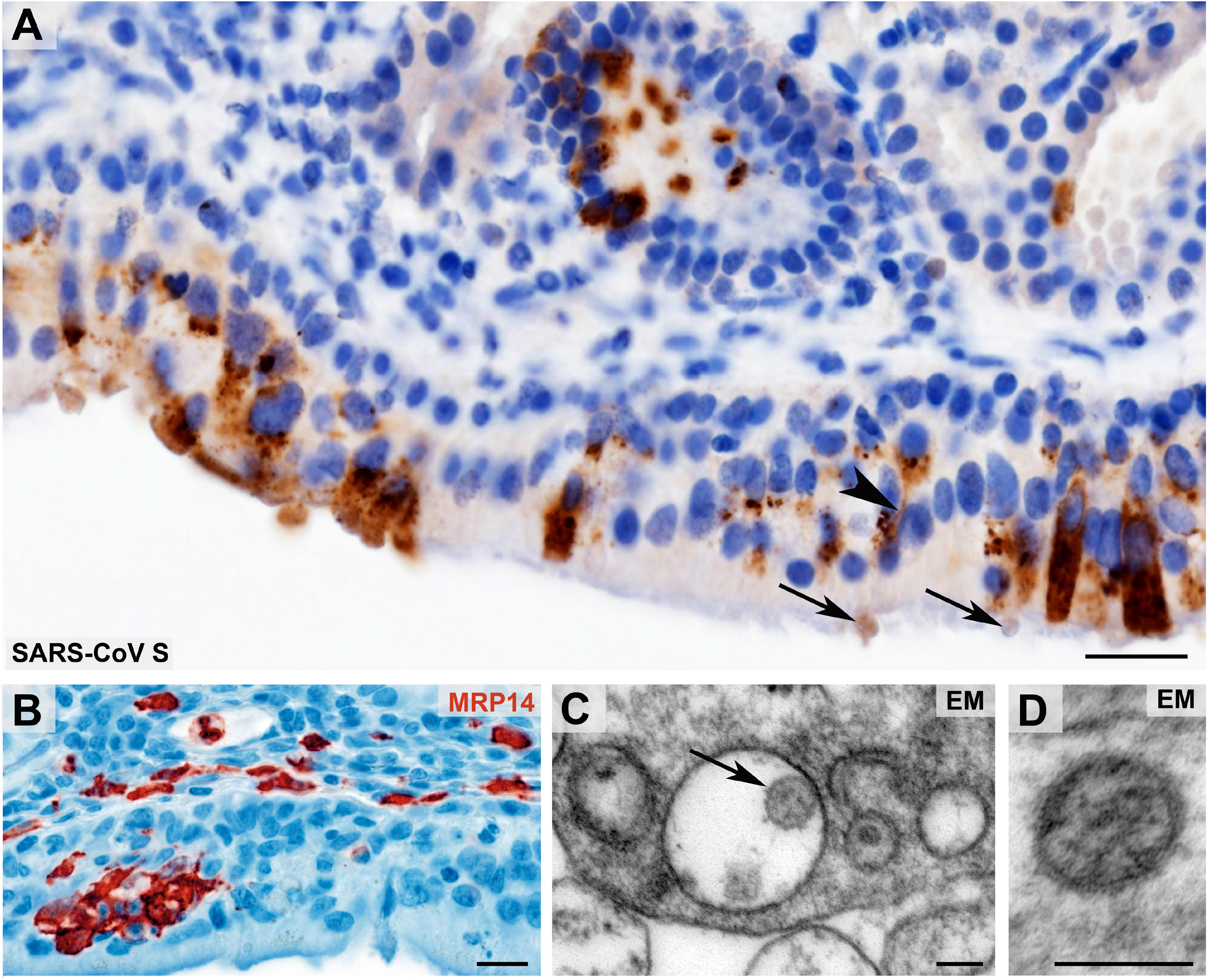
Morphological evidence of SARS-CoV presence and first innate immune cell response within the olfactory mucosa. Coronavirus antigen (A, SARS-CoV Spike Protein (SARS-CoV S), brown color) exhibits a cytoplasmic staining with perinuclear accentuation of infected mucosal (epithelial) cells and identifies SARS-CoV-positive dendrites (arrowhead) and vesicles at the dendrite tips (arrows) of the olfactory receptor neurons. Small clusters of infiltrating, early activated macrophages and granulocytes (MRP14, red color) in the olfactory epithelium upon SARS-CoV-2 infection (B). Ultrastructural images of two different examples of Coronavirus-like particles in the olfactory mucosa (C - D; arrow in C) fulfilling the criteria of size, shape, structural features (membrane, surface structures, electron dense material within the particle, resembling ribonucleoprotein) and localization (C, cytoplasmic localization within a membrane compartment, sometimes with typical attachment on the inner membrane surface as shown in this example; D, extracellular). Scale bars: A, B: 20 μm; C, D: 100 μm.

Additional support for SARS-CoV-2 persistence was provided by ultrastructural analyses of ultrathin sections. We found Coronavirus-like particles (Figure 3) – despite subtle ultrastructural differences compared to coronavirus derived from infected cell cultures providing a somewhat different milieu (Supplementary Figure 3) – fulfilling the criteria of size, shape, substructure (membrane, surface projections and internal electron dense material, resembling ribonucleoprotein) and intracellular localization of Coronavirus particles, while being clearly distinct from intrinsic cellular structures resembling Coronavirus particles^20–25^.

## Discussion

We provide first evidence that SARS-CoV-2 neuroinvasion occurs at the neuro-mucosal interface by transmucosal trespassing via regional nervous structures followed by a transport along the olfactory tract of the CNS, thus explaining some of the well-documented neurological symptoms in COVID-19 patients including alterations of smell and taste perception. Further studies are required to identify the precise cellular and molecular entry mechanism as well as receptors in the olfactory mucosa, where also non-neuronal pathways may play a role^26^. This will include a precise characterization of ACE2, TMPRSS2 and of CD147 expression, which have been implicated in enabling SARS-CoV-2 invasion in cells. Moreover, in line with recent clinical data demonstrating thromboembolic CNS events described in few patients^27^, we found in 13% of the 32 investigated cases also the histopathological correlate of microthrombosis and territorial brain infarcts.

The presence of genuine virus in the olfactory mucosa with its delicate olfactory and sensitive – partially axonally damaged (Supplementary Figure 1) nerves in conjunction with SARS-CoV-2 RNA manifestation preferentially in those neuroanatomical areas receiving olfactory tract projections (Figure 1) may speak in favor of SARS-CoV-2 neuroinvasion occurring via axonal transport. However, several other mechanisms or routes, including transsynaptic transfer across infected neurons, infection of vascular endothelium, or leukocyte migration across the blood-brain barrier (BBB), or combinations thereof, be it in addition or exclusive, cannot be excluded at present^14^.

The detection of – compared to values measured in the lower respiratory tract (Supplementary Table 1) – persistently high levels of SARS-CoV-2 RNA in the olfactory mucosa (124% as mean value compared to lower respiratory tract) up to 53 days after initial symptoms (Supplementary Table 1), including the detection of sgRNA suggests that olfactory mucosa remains a region of continuous SARS-CoV-2 replication and persistence, thus enabling a constant viral replenishment for the CNS. In line with our findings there is a comparable SARS-CoV-2 infection gradient from the nose to the lungs which is paralleled by expression of the receptor molecule ACE2^28^. Although this remains to be speculation and widespread dysregulation of homeostasis of cardiovascular, pulmonal and renal systems has to be regarded as the leading cause in fatal COVID-19 cases, previous findings of SARS-CoV infection and other coronaviruses in the nervous system^29^ as well as the herein described presence of SARS-CoV-2 RNA in the medulla oblongata comprising the primary respiratory and cardiovascular control center bring to mind the possibility that SARS-CoV2 infection, at least in some instances, can aggravate respiratory or cardiac problems – or even cause failure – in a CNS-mediated manner^6,30^.

Even when following distinct routes upon first CNS entry and – based on our findings – in the absence of clear signs of widespread distribution of SARS-CoV-2 in the CNS (i.e. no signs of meningitis/encephalitis in COVID-19 cases), it cannot be excluded that the virus may spread more widely to other brain regions, thus eventually contributing to a more severe or even chronic disease course, depending on various factors such as the time of virus persistence, viral load, and immune status, amongst others.

## Methods

### Study design

32 autopsy cases of either PCR-confirmed SARS-CoV-2 COVID-19 patients (N=29 of 32) or of patients clinically highly suggestive of COVID-19 (N=3 out of 32) were included. Autopsies were performed at the Department of Neuropathology and the Institute of Pathology, Charité – Universitätsmedizin Berlin (N=25 out of 32) including one referral case from the Institute of Pathology, DRK Kliniken Berlin, the Institutes of Pathology and of Neuropathology, University Medicine Göttingen (N=6 out of 32) and the Institute of Forensic Medicine Charité – Universitätsmedizin Berlin (N=1 out of 32). This study was approved by the Ethics Committee of the Charité (EA 1/144/13 and EA2/066/20) as well as by the Charité-BIH COVID-19 research board and was in compliance with the Declaration of Helsinki. In all deceased patients a whole-body autopsy was performed, which included a thorough histopathologic and molecular evaluation comprising virological assessment of SARS-CoV-2 RNA levels as indicated in Supplementary Table 1. Clinical records were assessed for pre-existing medical conditions and medications, current medical course, and ante-mortem diagnostic findings.

### SARS-CoV- and SARS-CoV-2-specific PCR including subgenomic RNA assessment

RNA was purified from ∼50 mg of homogenized tissue from all organs by using the MagNAPure 96 system and the MagNA Pure 96 DNA and Viral NA Large Volume Kit (Roche) following the manufacturer’s instructions.

Quantitative real-time PCR for SARS-CoV-2 was performed on RNA extracts with RT-qPCR targeting the SARS-CoV-2 E-gene. Quantification of viral RNA was done using photometrically quantified *in vitro* RNA transcripts as described previously^33^. Total DNA was measured in all extracts by using the Qubit dsDNA HS Assay kit (Thermo Fisher Scientific, Karlsruhe, Germany).

Detection of subgenomic RNA (sgRNA), as correlate of active virus replication in the tested tissue was done using oligonucleotides targeting the leader transcriptional regulatory sequence and region within the sgRNA coding for the SARS-CoV-2 E gene, as described previously^34^.

### Electron microscopy

Autopsy tissues were fixed with 2.5% glutaraldehyde in 0.1M sodium cacodylate buffer, postfixed with 1% osmium tetroxide in 0.05M sodium cacodylate, dehydrated using graded acetone series, then infiltrated and embedded in Renlam resin. *En bloc* staining with uranyl acetate and phosphotungstic acid was performed at the 70% acetone dehydration step. 500 nm semithin sections were cut using an ultramicrotome (Ultracut E, Reichert-Jung) and a histo jumbo diamond knife (Diatome), transferred onto glass slides, stretched at 120°C on a hot plate and stained with Toluidine blue at 80°C. 70 nm ultrathin sections were cut using the same ultramicrotome and an ultra 35° diamond knife (Diatome), stretched with xylene vapor, collected onto pioloform coated slot grids and then stained with lead citrate. Standard TEM was performed using a Zeiss 906 in conjunction with a 2k CCD camera (TRS). Large-scale digitization was performed using a Zeiss Gemini 300 field-emission scanning electron microscope in conjunction with a STEM-detector via Atlas 5 software at 4-6 nm pixel size. Regions of interest of the large-scale datasets were saved by annotation (“mapped”) and then recorded at very high resolution using 0.5-1 nm pixel size.

### Immunohistochemical procedures and stainings

Routine histological stainings (Hematoxylin and eosin (HE), Masson-Goldner, Periodic acid-Schiff reaction (PAS), and Toluidine blue) were performed according to standard procedures. Immunohistochemical stainings were either performed on a Benchmark XT autostainer (Ventana Medical Systems, Tuscon, AZ, USA) with standard antigen retrieval methods (CC1 buffer, pH8.0, Ventana Medical Systems, Tuscon, AZ, USA) or manually using 4-μm-thick FFPE tissue sections. The following primary antibodies were used: monoclonal mouse anti-S100 (DAKO Z0311, 1:3000), monoclonal mouse anti-AE1/AE3 (DAKO M3515, 1:200), monoclonal mouse anti-MRP14 (Acris, BM4026B, 1:500, pretreatment protease), monoclonal mouse anti-CD56 (Serotec, ERIC-1, 1:200), mouse monoclonal anti-SARS spike glycoprotein antibody (Abcam, ab272420, 1:100, pretreatment Citrate + MW) and polyclonal rabbit anti-OLIG2 (IBL, 18953, 1:150, pretreatment Tris-EDTA + MW). Briefly, primary antibodies were applied and developed either using the iVIEW DAB Detection Kit (Ventana Medical Systems) and the ultraView Universal Alkaline Phosphatase Red Detection Kit (Ventana Medical Systems) or by manual application of biotinylated secondary antibodies (Merck, RPN1001, RPN1004), peroxidase-conjugated avidin, and diaminobenzidine (DAB, Sigma, D5637) or 3-Amino-9-Ethylcarbazol (AEC). Sections were counterstained with hematoxylin, dehydrated in graded alcohol and xylene, mounted and coverslipped. IHC stained sections were evaluated by at least two board-certified neuropathologists with concurrence. For data handling of whole slides images an OME-TIFF workflow was used^35^.

## Acknowledgements

This work was supported by the Deutsche Forschungsgemeinschaft (DFG, German Research Foundation) under Germany’s Excellence Strategy – EXC-2049 – 390688087, as well as SFB TRR 167 and HE 3130/6-1 to F.L.H., SFB 958/Z02 to J.S., SFB TRR 130 to H.R., EXC 2067/1-390729940, SFB TRR 274 and STA 1389/5-1 to C.S., by the German Center for Neurodegenerative Diseases (DZNE) Berlin, and by the European Union (PHAGO, 115976; Innovative Medicines Initiative-2; FP7-PEOPLE-2012-ITN: NeuroKine). We are indebted to Francisca Egelhofer, Petra Matylewski, Kathrein Permien, Vera Wolf, Sandra Meier, René Müller, Uta Scheidt and Katja Schulz for excellent technical assistance and advice. We thank the Core Facility for Electron Microscopy of the Charité for support in acquisition of the data. The authors are most grateful to the patients and their relatives for consenting to autopsy and subsequent research, which were facilitated by the Biobank of the Department of Neuropathology – Universitätsmedizin Berlin, Germany. Furthermore, we thank the Charité foundation for financial support. Cartoon images were partially created with Biorender.com.

## Author Contributions

J.M., J.R., R.M., J.F., O.H., M.W., L.R., H.M., W.J.S-S., C.S., S.H., M.S.W., B.T., S.E., D.H., L.O., M.T, B.I.H., H.R., F.L.H. performed clinical workup and sections and/or histological analyses, Ca.D., H.H.G., M.L. and W.S. did ultrastructural analyses, J.S., S.B., Ch.D. and V.M.C. made viral RT-qPCR analyses. All authors contributed to the experiments and analyzed data; H.R. and F.L.H. designed and supervised the study; J.M., J.R., and Ca.D. prepared figures. All authors wrote, revised and approved the manuscript.

