## Supplementary material for "Olfactory transmucosal SARS-CoV-2 invasion as port of Central Nervous System entry in COVID-19 patients": Suppl. Figs 1-3 and Suppl Table 1

### Supplementary Figures 1 – 3 incl. Fig. legends

**Olfactory transmucosal SARS-CoV-2 invasion as port of Central Nervous System entry in COVID-19 patients by Meinhardt et al.**

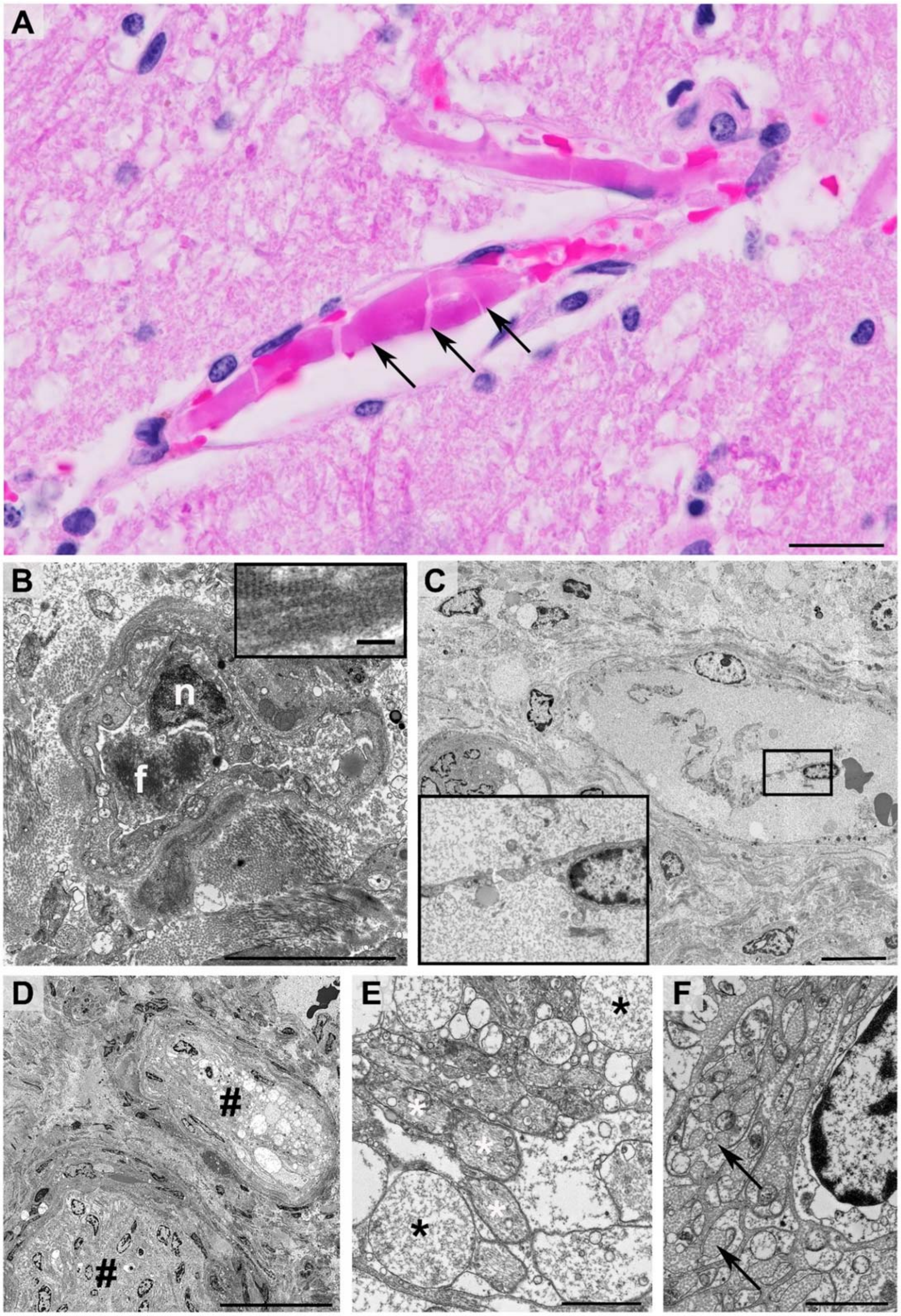


**Supplementary Figure 1: (Micro)thromboembolic events in thalamus and olfactory mucosa of patients dying of COVID-19**

Hematoxylin-Eosin (HE)-stained formalin-fixed paraffin-embedded (FFPE) section (A) illustrating the thalamus of a COVID-19 autopsy case (patient 26). Several small vessels exhibited fresh thrombi (A, pink color, arrows) resulting in a large infarct of surrounding CNS tissue characterized by a substantial reduction of detectable neuronal and glial nuclei, oedema and vacuolation. Ultrastructural aspect of the olfactory mucosa (patient 8) demonstrating vessels, some of which were filled with heterogeneous electron dense filamentous material resembling fibrin (B with f: fibrin depicted in transverse orientation, n: endothelial cell nucleus), which revealed its characteristic periodic pattern when exhibited in longitudinal orientation (insert in B). Vascular/endothelial pathology included endothelial detachment (C, inset showing an ellipse-shaped nucleus and flat cytoplasmic extensions demonstrating a polar orientation characteristic for endothelial cells). Nerve fascicles in the lamina propria (#) exhibiting focal vacuolation (upper nerve fascicle, magnified in E) as signs of acute damage (D). Damaged nerve fascicle with degenerating axons (black asterisks in E) and remaining intact axons (white asterisks). (F) Distinct collagen deposition (arrows), i.e. collagen pockets, corresponding to loss of unmyelinated axons in damaged nerve fascicles (magnification of the lower nerve fascicle shown in D). Scale bars: A: 50 µm; B: 5 µm (inset in B: 100 nm); C: 10 µm; D: 30 µm; E, F: 2 µm.

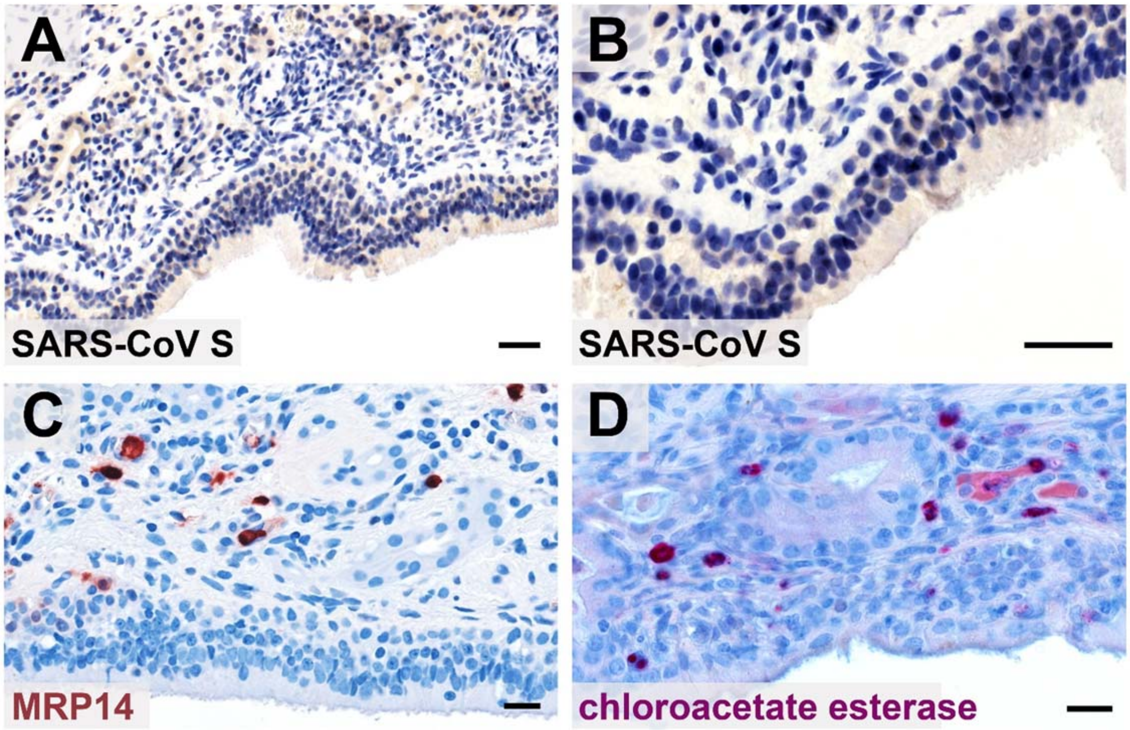


**Supplementary Figure 2: MRP14 and SARS-CoV s control stainings**

Immunohistochemical staining of the olfactory mucosa from an age-matched non-COVID-19 patient shows absence of SARS-CoV spike protein signal (A, B) and no MRP14 expressing cells in the olfactory mucosa (C). Only in the submucosa, few MRP14-positive signals can be detected. As MRP14 is expressed by early activated macrophages and granulocytes, enzymatic chloroacetate esterase staining considered specific for cells of granulocytic lineage on serial sections of the MRP14-stained tissue shown in Figure 3B revealed only single granulocytes in the olfactory mucosa (D) corroborating our interpretation that the majority of MRP14-expressing cells in COVID-19 patients (shown in Figure 3B) are early activated intramucosal macrophages. Scale bars A - D: 20 µm.

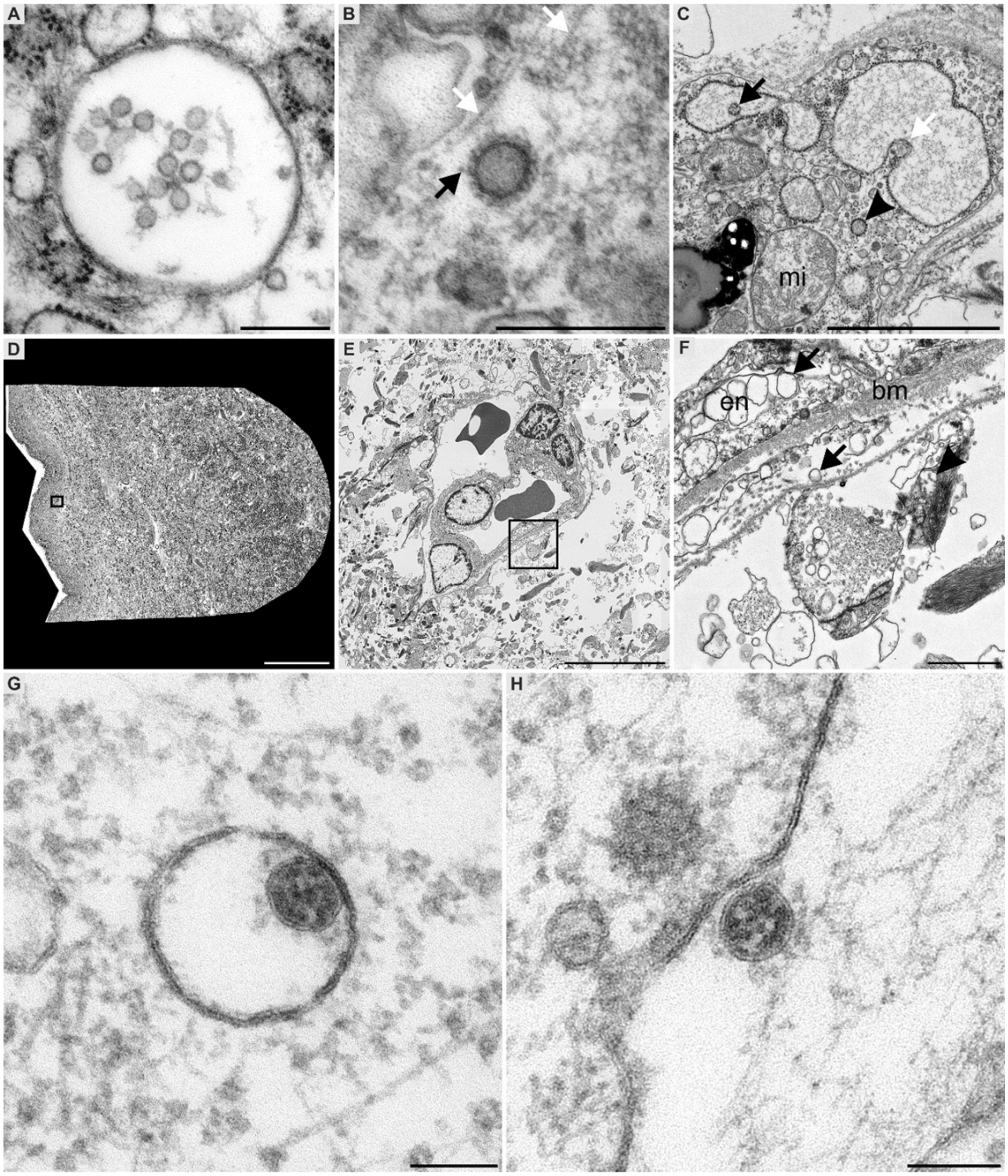


**Supplementary Figure 3: Morphological mimics and autolysis as ultrastructural pitfalls in coronavirus identification in autopsy tissues and SARS-CoV-2 ultrastructural presentation under cell culture conditions**

SARS-CoV-2 PCR-positive COVID-19 lung autopsy samples (A – C; A+B: patient 6; C: patient 7) and medulla oblongata (D – F; patient 6) with low numbers of RNA copies. Subcellular structures such as multi-vesicular bodies (MVBs; A) and coated vesicles (B) are easily mistaken as coronaviruses[21]. However, in contrast to coronavirus, MVBs lack characteristic substructure represented by electron dense ribonucleoprotein complexes and delicate surface projections. Additionally, size and position of the vesicles are different in coronavirus, e.g. coated vesicles (B) are not inside a membrane compartment (note the cytoplasmic microtubules indicated by white arrows), lack the substructure, and show too pronounced and regular surface projections (black arrow). Invaginations (C, white arrow) of e.g. rough endoplasmic reticulum cisternae (rER, pneumocyte) may also mimic coronavirus in some cross-sectional orientations (compare to vesicular structure; black arrow), due to high electron density and granular appearance of ribosomes, resembling ribonucleoprotein complex (RNP); note cytoplasmic vesicle, most likely of the rER, with granular appearance (C, arrowhead). Large-scale digitization of the entire ultrathin section of post-mortem medulla oblongata tissue (D - F) at a pixel size of 4 nm. (E) and (F) showing digitally magnified areas of the insets in (D) and (E), demonstrating post-mortem-related changes e.g. vacuolation (E - F, arrows in F), swollen mitochondria (arrowhead in F) blurring coronavirus detection compared to freshly prepared human or laboratory animal specimens or cell culture. Ultrastructure of SARS-CoV-2 (isolate Italy-INMI1) virus particles in Vero E6 cells (G, H: 24 h post infection) demonstrate the intracellular localization within a membrane compartment (G) with a typical attachment to the inner membrane surface. Note the preservation of surface projections which is due to the application of a particular *en bloc* staining (tannic acid and uranyl acetate)[32]. Electron dense material, likely representing RNP, can be detected directly underneath the envelope membrane and within the particle lumen. Extracellular virus particle with similar structural appearance and attachment to the apical plasma membrane (H). Scale bars: A, B: 250 nm; C: 2000 nm; D: 250 µm; E: 10 µm; F: 1000 µm; G, H: 100 nm.

| | | | | | | Pre-existing Conditions / Concomitant Diseases | | | | | | log10SARS-COV-2-RNA copies/10,000 cells | | | | | | | | | |
|---|---|---|---|---|---|---|---|---|---|---|---|---|---|---|---|---|---|---|---|---|---|
| **Patient No.** | **Gender** | **Age [years]** | **Postmortem interval [hours ]** | **SARS-Cov-2 qPCR ante-mortem** | **Duration of illness [days]** | Cardiovascular | Respiratory | Gastrointestinal | Urinary | Cerebral | Miscellaneous | Carotis | Cornea | Conjunctiva | Uvula | Olfactory mucosa | Olfactory bulb | Trigeminal ganglion | Medulla oblongata | Cerebellum | olfactory mucosa normalized to highest lung value |
| **1** | M | 71 | 112 | Positive | 34 | Arterial hypertension | Bronchial asthma | _ | _ | _ | _ | N/A | N/A | N/A | N/A | 2,01 | 0 | N/A | 0 | N/A | 0,81 |
| **2** | F | 79 | 148 | Positive | 11 | Arterial hypertension, coronary heart disease | _ | _ | Stage 3 chronic kidney disease | Dementia | Obstructive sleep apnoea | N/A | N/A | N/A | 1,84 | 5,06* | 3,26 | N/A | 0 | N/A | 0,86 |
| **3** | F | 90 | 106 | Positive | 9 | Arterial hypertension, coronary heart disease, atrial fibrillation | COPD, stage 2 (Gold classification) | Liver cirrhosis | Stage 3 chronic kidney disease | _ | _ | N/A | N/A | N/A | 3,22 | 7,47* | 3,1 | 0 | 1,21 | N/A | 1,21 |
| **4** | M | 79 | 56 | Positive | 28 | _ | _ | _ | _ | _ | Right hemiplegia since childhood | N/A | N/A | N/A | 0 | 0 | 0 | 0 | 0 | 0 | |
| **5** | F | 79 | 64 | Positive | 16 | Arterial hypertension, coronary heart disease | _ | _ | _ | Left middle cerebral artery stroke, right hemiataxia (2019) | Type 2 diabetes mellitus, Osteoporosis, Hyperlipidemia | N/A | N/A | N/A | 0 | 0 | 0 | 0 | 2,74 | 0 | |
| **6** | M | 68 | 30 | Positive | 35 | _ | _ | _ | _ | _ | _ | N/A | N/A | N/A | 0 | 3,29 | 0 | 1,67 | 2,03 | 0 | 1,90 |
| **7** | M | 61 | 10 | Positive | 21 | Arterial hypertension, coronary heart disease | Heavy smoker | Cholecystectomy | _ | _ | Class 1 Obesity | N/A | N/A | N/A | 0 | 0 | 0 | 0 | 0 | 0 | |
| **8** | F | 68 | 16 | Positive | 37 | Arterial hypertension, coronary heart disease | COPD, stage 2 (Gold classification) | _ | _ | _ | Class 1 Obesity, Hypothyroidism | N/A | 2,42 | 0 | 0 | 2,39 | 0 | 0 | 0 | 0 | 1,93 |
| **9** | M | 76 | 30 | Positive | N/A | Arterial hypertension, ischemic cardiomyopathy, myocardial infarction (2008) | Heavy smoker | _ | _ | _ | _ | N/A | 0 | 0 | 1,29 | 5,57* | 0 | 2,05 | 0 | 0 | 0,85 |
| **10** | F | 56 | 38 | Positive | 26 | Arterial hypertension | _ | _ | _ | _ | Type 2 diabetes mellitus, Class 1 obesity, breast cancer (2007) | N/A | N/A | 0 | 0 | 0 | 0 | 0 | 0 | 1,94 | |
| **11** | M | 92 | 72 | Positive | 30 | Intermittent atrial fibrillation | _ | _ | _ | Dementia | _ | N/A | N/A | 0 | 0 | 0 | N/A | 0 | 1,46 | 0 | |
| **12** | F | 67 | 48 | Positive | 26 | Arterial hypertension | _ | Sigma diverticulosis | _ | Right frontal intracerebral hemorrhage with infarction (2009), left-sided hemiparesis, VP-Shunt (2010) | Rheumatoid arthritis | 0 | N/A | 0 | N/A | N/A | 0 | 0 | 0 | 0 | |
| **13** | M | 62 | 144 | Positive | 33 | Arterial hypertension | Heavy smoker | _ | _ | Vascular dementia, left-sided watershed stroke (2016), right-sided hemiparesis | Hyperlipo-proteinemia | N/A | 0 | 0 | 0 | 1,32 | 0 | 0 | 0 | 0 | 0,78 |
| **14** | F | 78 | 192 | n.d., clinical presentation suggestive of COVID-19 | N/A | Arterial hypertension, atrial fibrillation, aortic/mitral valve insufficiency | Small cell lung cancer | _ | _ | _ | _ | 0 | N/A | 0 | N/A | N/A | 0 | N/A | 0 | 0 | |
| **15** | M | 81 | 82 | Positive | 4 | Arterial hypertension, atrial fibrillation, abdominal aortic aneurysm | _ | _ | _ | Ischemic stroke with left-sided hemiparesis | Type 2 diabetes mellitus, osteoporosis, hyperthyroidism | N/A | 0 | N/A | 6,4* | 7,98* | 2,89 | 2,17 | N/A | 0 | 1,15 |

| | | | | | | | | | | | | | | | | | | | | | |
|---|---|---|---|---|---|---|---|---|---|---|---|---|---|---|---|---|---|---|---|---|---|
| **16** | M | 74 | 40 | Positive | 53 | Arterial hypertension | _ | _ | Nephrolithiasis | _ | Class 2 obesity | 0 | 0 | 0 | 0 | 3,21 | 0 | 0 | 0 | 0 | 1,48 |
| **17** | M | 70 | 116 | Positive | 48 | _ | _ | _ | Bladder cancer (1999; treated curatively) | _ | _ | 0 | 3,16 | 2,12 | 0 | 3,04 | 0 | 0 | 0 | 0 | 1,27 |
| **18** | F | 30 | 38 | n.d., clinical presentation suggestive of COVID-19 | 21 | _ | CMV reactivation + toxoplasmosis with pulmonary involvement | Intestinal acute graft versus host disease (GvHD) | _ | Cerebral toxoplasmosis | Allogenic stem cell transplantation (2019), Ewing sarcoma (2013) | 0 | 0 | 0,8 | 0 | 0 | 0 | 0 | 0 | 0 | |
| **19** | M | 77 | 53 | Positive | 44 | Arterial hypertension | Pulmonary emphysema | _ | _ | _ | _ | 0 | 0 | 0 | 0 | 0 | 0 | 0 | 0 | 0 | |
| **20** | M | 57 | 46 | Positive | 36 | _ | _ | _ | _ | _ | _ | 0 | 0 | 0 | 0 | 1,66 | 0 | 0 | 0 | 1,51 | |
| **21** | M | 79 | 168 | Positive | N/A | _ | Heavy smoker | Hepatic steatosis, cholelithiasis | Kidney cysts | _ | Psoriasis, gonarthrosis | N/A | N/A | N/A | 1,29 | 0 | 0 | N/A | 0 | 0 | |
| **22** | M | 69 | 94 | Positive | 39 | Arterial hypertension | _ | _ | _ | _ | Polyneuropathy, alcohol abuse (abstinence for 18 years) | 0 | 2,6 | 0 | 0 | 1,9 | 0 | 0 | 0 | 0 | 1,19 |
| **23** | M | 81 | 28 | n.d., clinical presentation suggestive of COVID-19 | N/A | _ | _ | _ | _ | Acute pontine infarction with dysarthria and dysphagia (03/2020) | Type 2 diabetes mellitus | 0 | 0 | 0 | 0 | 0 | 0 | 0 | 0 | 0 | |
| **24** | M | 74 | 216 | Positive | 35 | Coronary heart disease, severe aortic valve stenosis, biologic aortic valve prothesis | _ | _ | _ | _ | _ | 0 | 0 | 0 | N/A | 2,54 | 0 | 0 | 0 | 0 | 1,41 |
| **25** | M | 62 | 72 | Positive | 15 | Arterial hypertension | _ | Upper gastrointestinal bleeding | _ | _ | Obstructive sleep apnoea, type 2 diabetes mellitus, | N/A | N/A | N/A | N/A | N/A | N/A | N/A | N/A | N/A | |
| **26** | F | 45 | 48 | Positive | 20 | Arterial hypertension | Bronchial asthma | _ | _ | _ | Alcohol abuse | N/A | N/A | N/A | N/A | N/A | N/A | N/A | N/A | N/A | |
| **27** | M | 71 | 11 | Positive | N/A | _ | _ | _ | _ | Parkinson's disease | _ | N/A | N/A | N/A | N/A | N/A | N/A | N/A | N/A | N/A | |
| **28** | M | 80 | 72 | Positive | 30 | Arterial hypertension, atrial fibrillation, endocarditis (2014) | _ | _ | Stage 4 chronic kidney disease | _ | Granulomatosis with polyangiitis, anemia, polyneuropathy, osteoporosis | N/A | N/A | N/A | N/A | N/A | N/A | N/A | N/A | N/A | |
| **29** | F | 72 | <24 | Positive | N/A | _ | _ | _ | _ | _ | _ | N/A | N/A | N/A | N/A | N/A | N/A | N/A | N/A | N/A | |
| **30** | F | 98 | 96 | Positive | 4 | Arterial hypertension, atrial fibrillation, bradycardia | _ | _ | _ | Left middle cerebral artery stroke | _ | N/A | N/A | N/A | N/A | N/A | N/A | N/A | N/A | N/A | |
| **31** | M | 68 | 20 | Positive | 35 | _ | _ | _ | _ | _ | _ | N/A | N/A | N/A | N/A | N/A | N/A | N/A | N/A | N/A | |
| **32** | M | 68 | 24 | Positive | 58 | Arterial hypertension, atrial fibrillation, coronary heart disease, myocardial infarction | COPD | Acute pancreatitis | Stage 2 chronic kidney disease | Basilaris thrombosis with mechanical thrombectomy | Bullous pemphigoid, type 2 diabetes mellitus | N/A | N/A | N/A | N/A | N/A | N/A | N/A | N/A | N/A | |

*** subgenomic RNA (sgRNA) positive**
